# Response to Tabin *et al* “Concerns about ancient DNA sequences reported from a Late Pleistocene individual from Southeast Asia”

**DOI:** 10.1101/2025.03.29.646132

**Authors:** Xiao-Ming Zhang, Yao-Xi He, Xue-Ping Ji, Bing Su

**Affiliations:** Key Laboratory of Genetic Evolution & Animal Models, Kunming Institute of Zoology, Chinese Academy of Sciences, Kunming 650201, China; Kunming Natural History Museum of Zoology, Kunming Institute of Zoology, Chinese Academy of Sciences, Kunming 650223, China; Yunnan Key Laboratory of Integrative Anthropology, Kunming Institute of Zoology, Chinese Academy of Sciences, Kunming 650201, China

## Abstract

Tabin *et al^1^* suspect a high error rate and abnormal error content in the MZR genome data from our published study^2^, from which they raised concerns about the reliability and useability of our published sequences. Given the poor environmental conditions (such as warm climate and acidic soil in the low latitude area of Southwest China), as well as the non-ideal fossil material (cranium) for DNA extraction, we argue that a relatively high level of aDNA damage, as well as possible artefacts from extraction, library construction and sequencing better explain the observed pattern by Tabin *et al* in MZR rather than modern DNA contamination. Particularly, we think the mutation motif of the MZR mtDNA, derived from our careful manual check, should be reliable. In addition, we provide additional analyses showing how we minimize the effect of aDNA damage in population analyses.

## Main text

Tabin *et al*^*1*^ observed lots of mismatch sites in the MZR mtDNA sequence when compared to the rCRS reference. However, based on the pattern of sequence mismatches, we argue that the great majority of these mismatches exhibit high rates of allele consistency, and they do not influence mtDNA haplogroup determination. In addition, we provide lines of evidences to exclude the possibility that these low counts of derived allele reads were caused by modern DNA contamination.

There are in general only 40-50 mutations in the mtDNA genome of a modern human individual. Consequently, as more than 3,000 mismatches detected in MZR mtDNA genome by Tabin *et al*^*1*^, at least 60 individuals would be presumably involved in the proposed contamination, which is unlikely because only a few people were able to have contact with the MZR sample, and we had conducted a thorough surface cleaning before bone sample collection and DNA extraction. Second, among the 3,058 mismatch sites, we found 2,284 of them (75%) being transversion mismatches (Table S1). In the human mtDNA population data, there are only 2-3 transversion mutations in one individual (the mutations list along the mtDNA tree Build in http://phylotree.org/). Hence, the frequently observed transversions are better explained by DNA damage or sequencing error, and these mismatches should not be counted when estimating the rate of modern human DNA contamination. Third, we sequenced the mtDNA genome of the person who performed all laboratory works of MZR, and his mtDNA belongs to the D4b2b4 lineage. The mutation motif of D4b2b4 is largely different from the MZR lineage (M9). Lastly, we searched the frequencies of the 3,058 mismatch sites in a large set of Han Chinese (21,927 sequences) and East Asians (27,850 sequences), and we found 73.62% of them are totally absent in the Han Chinese set, and 21.53% of them are rare mutations in Han Chinese (1%-3%) (Figure 1A; Table S1), implying a very low possibility of modern DNA contamination. To verify the defined M9 lineage of MZR mtDNA, using the untrimmed mapping data, we took a more conservative approach by only including the damage-enriched reads filtered by *PMDtools* (threshold=3) (Figure S1) so that the suspected modern DNA contamination can be faithfully removed. Based on this mtDNA sequence data with careful manual check, we confirmed the assigned M9 haplogroup of MZR mtDNA (Table S2).

**Figure 1.**
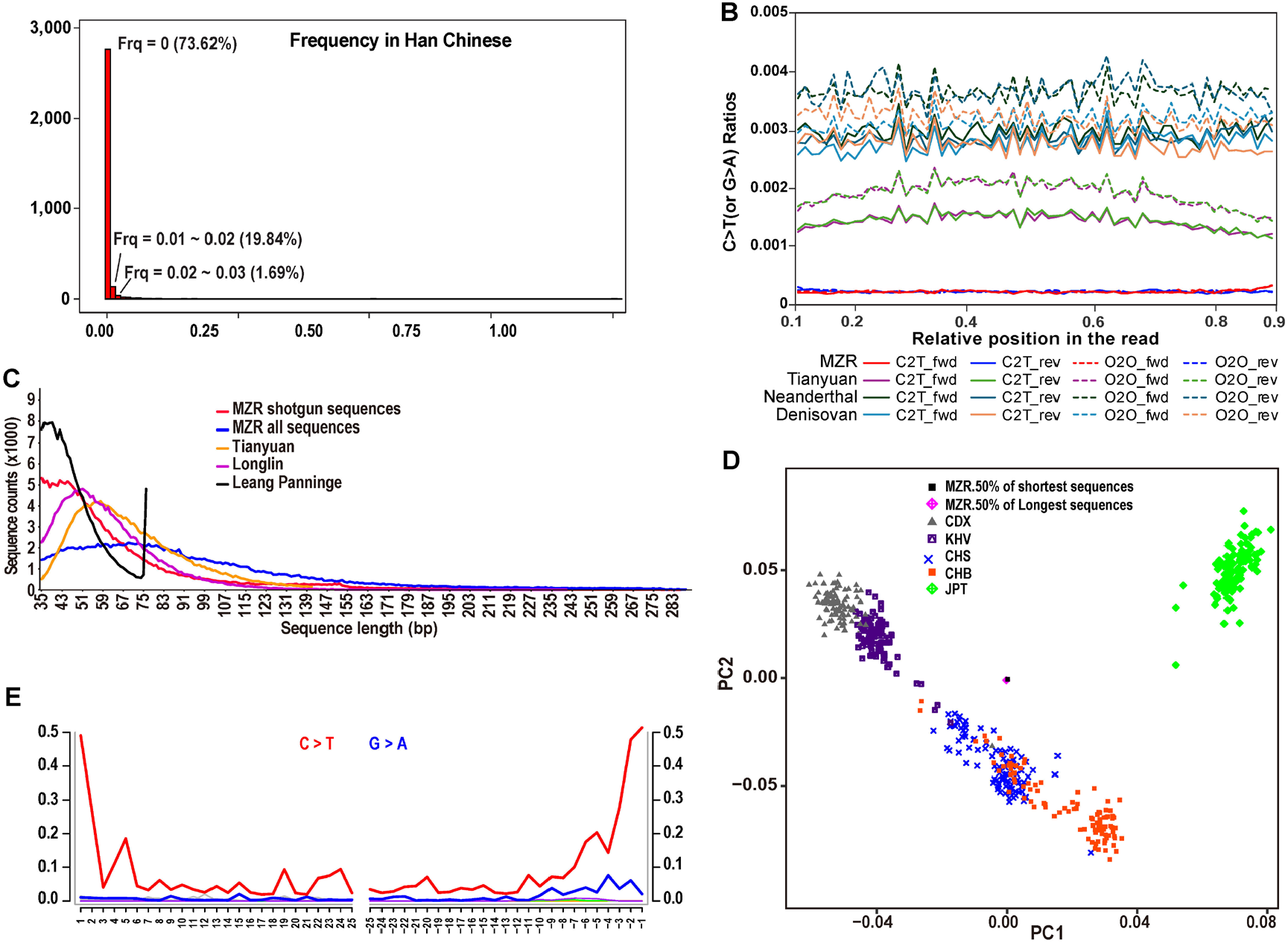
General evaluation of the MZR data. (A) The frequency distribution of the 3,058 MZR mtDNA mismatches in 21,927 Han Chinese mtDNA sequences. (B) Comparison of C>T (forward strand) and G>A (reverse strand) substitution ratios along the sequence reads between MZR, NEA (Altai-Neanderthal), DEN (Denisovan) and TY (Tianyuan). For comparison, we also plotted the ratios of the non-C>T/G>A substitutions. The non-standard substitutions (i.e. the non-C>T/G>A substitutions) along the sequence reads of MZR are shown by the light-colored curves. (C) The sequence length distribution in MZR (shotgun sequences and all sequences sets) and other published Late Pleistocene and early Holocene individuals from East Asia and Southeast Asia, including Tianyuan, Longlin and Leang Panninge. All the samples were randomly sampling ~187,543 reads which close to the fewest set (MZR shotgun sequences). (D) Using 2.2 million SNPs (1,240K+870K) set, the PCA plot showing the relationship among MZR (50% shortest sequences and 50% longest sequences sets) and present-day East Asians from the 1,000 Genomes Project3. We project the MZR data onto the first and second PC space established using the modern samples. (E) The pattern of terminal damages using the pooled single-strand libraries sequences with the PMDtools-filtered reads (threshold=3).

Based on the aforementioned observation of mismatches in MZR’s mtDNA, Tabin *et al*^*1*^ suspected the plausibility of the nuclear genome of MZR. However, for the nuclear genome data of MZR, as described in the original paper^2^, by strictly trimming many base pairs (1 to 2 bp at the 5’ end and 2 to 17 bp at the 3’) from the sequences, we thoroughly eliminated the DNA-damage-caused mismatches of MZR, and consequently, the calculated mutation ratios of MZR is comparable with the published ancient individuals (Figure 1B). The high consistency of the mtDNA lineage diagnostic SNPs also reflects a low level of DNA-damage-caused mismatches remained after the sequence ends trimming and filtering (Table S2, and Data S1D in the original paper). In addition, as expected, the DNA damage-caused transversion sites tend to show multiple alleles when merging with the published reference data, and they were completely removed from the downstream analyses. Indeed, we observed very similar proportions of different mutation types of the MZR genome when compared to the 1,240K SNP sets of Denisovan, Altai Neanderthal and Tianyuan (Figure 2).

**Figure 2.**
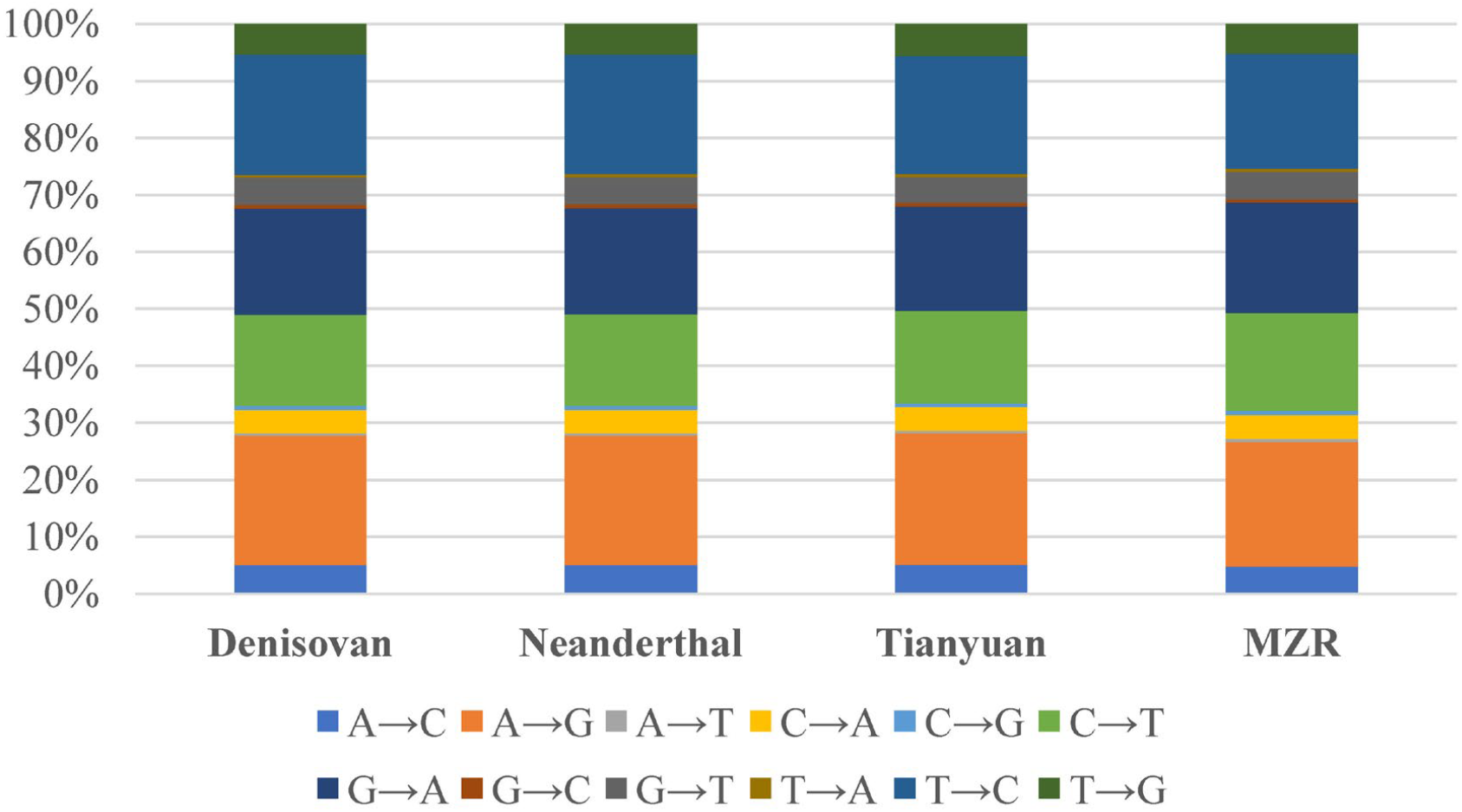
Mutations type proportion comparison in the genome of MZR with the published 1,240K SNP sets of Denisovan, Altai Neanderthal and Tianyuan (Table S3).

It is known that modern DNA contaminants tend to have longer fragmental length than those of ancient, endogenous DNA^4^. Regarding the sequence length of MZR, we summarized the length distribution of the representative Late Pleistocene and early Holocene individuals from East Asia and Southeast Asia, including MZR, Tianyuan^5^, Longlin^6^ and Leang Panninge^7^, and we did not see much length difference among the published data (Figure 1C), neither did we see unusual sequence length among the MZR libraries (Figure S2). For example, the average length of the MZR shotgun sequences is 64 bp, shorter than the sequence length of Tianyuan (70bp). In addition, the average fragmental length of the damage-enriched reads is 92bp, similar with the average length (93bp) of all the generated MZR reads (Figure S3), suggesting that the long fragment length is unlikely due to modern DNA contamination.

The baits length used for MZR is ~150 bp^8,9^. The myBaits kit from MYcroarray is even longer than the currently used version (the myBaits kit from Arbor Biosciences), which is also longer than the commonly used 1,240K baits (80 bp)^10^. Based on the baits length distribution detected using the High Sensitivity DNA Kit of Agilent Bioanalyzer 2100 (www.agilent.com/chem/labonachip) (Figure S4), we have confirmed that the used myBaits probe length from MYcroarray is ~150 bp, which is probably responsible for the relatively longer length of the MZR sequences (average 93 bp), rather than the suspected contamination.

Additionally, the PCA map indicates almost the same positions of the two sets of “MZR 50% of shortest sequences” and “MZR 50% of longest sequences” (Figure 1D), so are the two sets of “MZR original sequences” and “MZR damage enriched sequences” (Figure S5), reflecting a negligible difference among the different MZR data sets, again indicating a low level of DNA contamination in the MZR genome data.

Furthermore, for the DNA damage pattern of MZR, we did see a 3’-biased damage pattern when including both the data from single-stranded and double-stranded libraries, which might be due to biochemical processing during library construction or bioinformatics issue. This bias was also observed in several previous aDNA studies^11-13^. However, when we use the *PMDtools*^14^*-*filtered reads (threshold=3) from the single-stranded libraries for estimating autosomal contamination, we saw clear damage patterns at both the 5’ and 3’ ends (Figure 1E), indicating that the MZR data indeed contains the typically damaged DNA fragments. Using the *AuthentiCT*^15^ tool, we estimated the level of autosomal contamination based on the *PMDtools*-filtered reads, and the rate is very small (0.10%) for the single-stranded libraries of MZR. We also used the *conditional C2T mismatch*-based method^16^, and the estimated the level of contamination is also very low (0.72%) as already indicated in our original publication^2^. Finally, By generating a new 8.65 million SNPs dataset, Tabin *et al*^1^ observed that *D*(MZR, AR19K; Ancient USA Anzick, Ancient Cameroon) is Z = −3.6 standard errors below zero, which is inconsistent with our calculation using different number of SNPs in our original paper^2^. Based on this point, Tabin *et al*^1^ questioned our inference that during the Late Pleistocene, there was an express northward expansion of AMHs starting in southern East Asia through the coastal line of China, and eventually crossing the Bering Strait and reaching the Americas. Firstly, in the *D-stat* test, we found that when using different numbers of SNPs, the test could show inconsistency, e.g. for the datasets of 1,240K SNPs, 1,240K+870K SNPs, dbSNP147 (136,968,574 SNPs) and dbSNP147 (filter MAF=1%), we noticed that some *D-stat* tests could generate different D-values and Z-scores (Table S4). Secondly, using the new V62.0 of Allen Ancient DNA Resource (AADR), we comprehensively performed *D-stat* tests among the FirstAmericans (merged with 24 samples of ≥ 7kya), Paleo-Siberians (≥ 7kya), China_NEastAsia_Coastal_EN (merged with 6 samples of ≥ 7kya), China_SEastAsia_Coastal_EN (merged with 7 samples of ≥ 7kya), Japan_Shikoku_InitialJomon (8.8kya), and MZR (14.0kya). From the results, we observed increasing affinity to FirstAmericans from south to north among MZR, China_SEastAsia_Coastal_EN, China_NEastAsia_Coastal_EN, Russia_Boisman_MN and Russia_DevilsCave_N (Figure 3, Table S5). These results support the proposed early south to north human migration along coastal East Asia that eventually reached the Americas by way of Bering Strait. Thirdly, our previous study on modern human Y-chromosome DNA also supports the proposed migration route. The geographic distribution of the Y-chromosome C-M130 lineage suggests a prehistoric migration from south to north in mainland East Asia likely following the coastline, reaching Siberia and finally made its way to the Americas^17^. Fourthly, the coastal spatio-temporal distributions of mitochondrial DNA D4h lineage in eastern China and Thailand (D4h3b), Japanese archipelago (D4h1a and D4h2) and the Americas (D4h3a), in combination with the Paleolithic archaeological similarities among Northern China, Japan and the Americas, provides further support to the coastal dispersal scenario of early Native Americans^18^.

**Figure 3.**
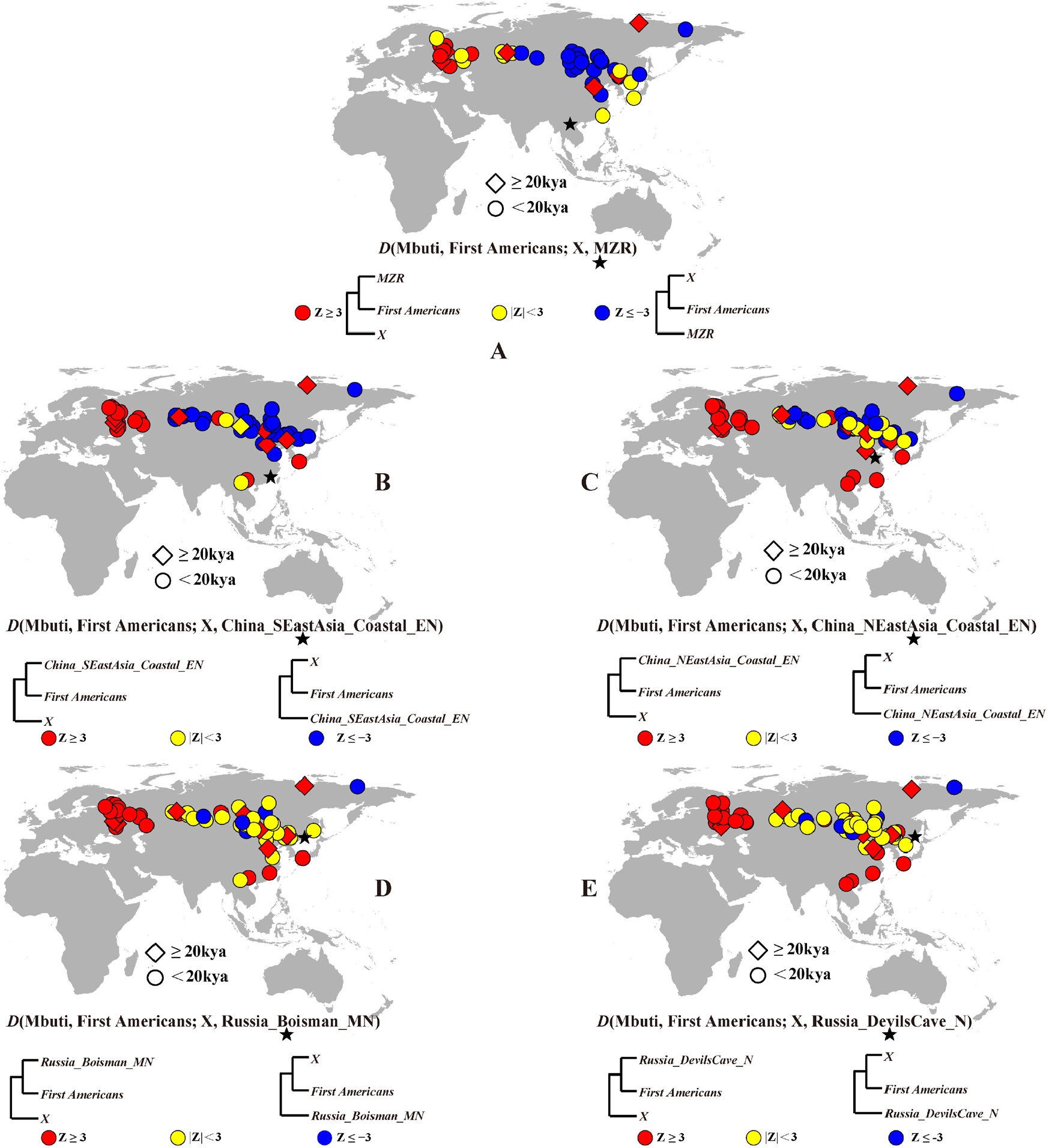
The test results of D(Mbuti, First Americans; X, Y), comparing the affinity degree to First Americans among Paleo-Siberians, MZR, China_SEastAsia_Coastal_EN, China_NEastAsia_ Coastal_EN, Russia_Boisman_MN, Russia_DevilsCave_N and Japan_Shikoku_InitialJomon. The results of pairwise SNPs number<20,000 were excluded in the analyses (Table S5).

Lastly, it should be noted that if we apply a stringent filtering criterion and only keep sequences showing characteristic of ancient DNA damage, the dataset would become very limited because a large portion of the MZR data was from the libraries with repaired-end damages by applying uracil-DNA-glycosylase (UDG) treatment. Fortunately, we have obtained additional MZR fossil specimens, including teeth, mandible and femur^19-21^, which are potentially better for aDNA preservation compared to the published skull aDNA data, and we are making efforts in obtaining more authentic genomic data by taking advantage of the 1,240K array capture^11^ (using the custom product by Twist Biosciences) so that the MZR genome data is more comparable to the published aDNA data for further in-depth population analysis.

## SUPPLEMENTAL INFORMATION

Supplemental information contains five figures, five tables, and supplemental methods.

## ACKNOWLEDGEMENTS

We thank Dr. Daniel Tabin and his colleagues from Dr. David Reich’s lab for their collegial interactions throughout the discussion. This work was funded by the National Natural Science Foundation of China (NSFC) (T2222030 and U23A20161), and Yunnan provincial “Ten Thousand Talents Plan-Youth Top Talent” project (YNWR-QNBJ-2019-160) to X.M Zhang.

## AUTHOR CONTRIBUTIONS

X.M Zhang and Y.X He performed analyses, X.M Zhang and B. Su wrote the letter. All authors edited and approved the letter.

## SUPPLEMENTAL INFORMATION

**Figure S1.**
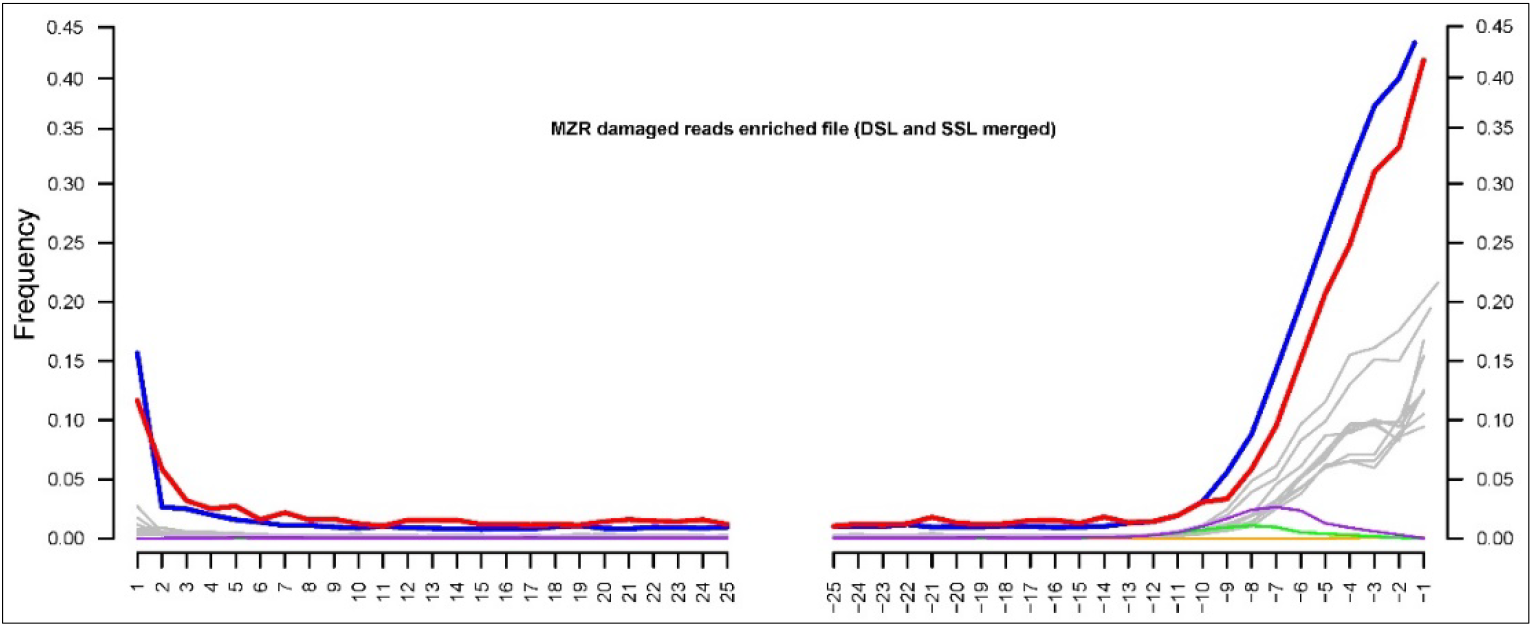
The MZR terminal damage profile after enriching the damaged reads using *PMDtools* (the DSL and SSL data were merged together). Using the un-trimmed raw sequence data, we re-mapped the MZR data. For the damage enriched sequence file, to eliminate the effect of DNA damage, before downstream SNPs calling (1,240K+870K SNPs set and dbSNP147 set), we trimmed 3bp and 10bp from the 5’ and 3’ ends, respectively.

**Figure S2.**
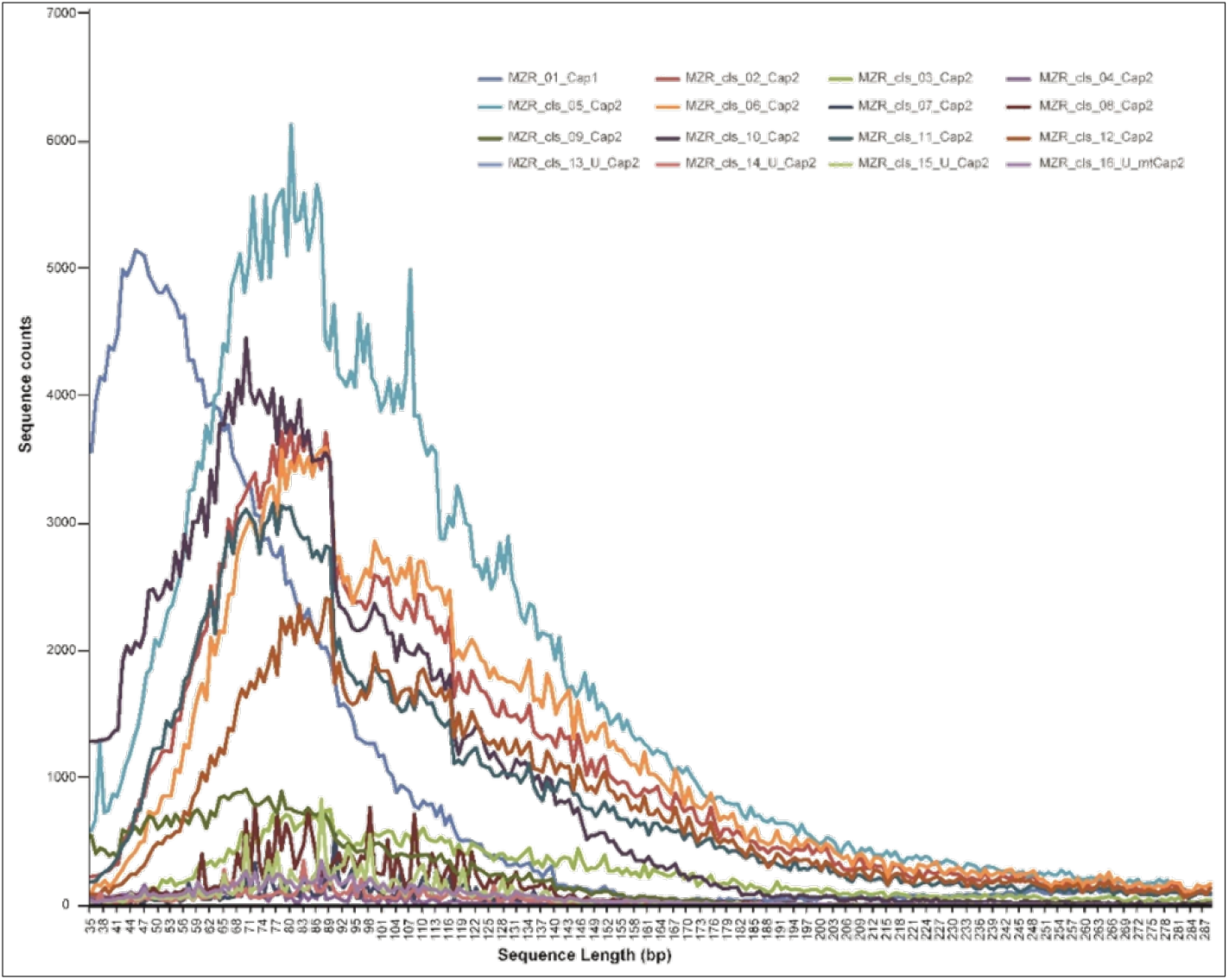
The sequence length distribution of the captured MZR libraries. The library symbols are the same as in Supplementary Table A in our original paper.

**Figure S3.**
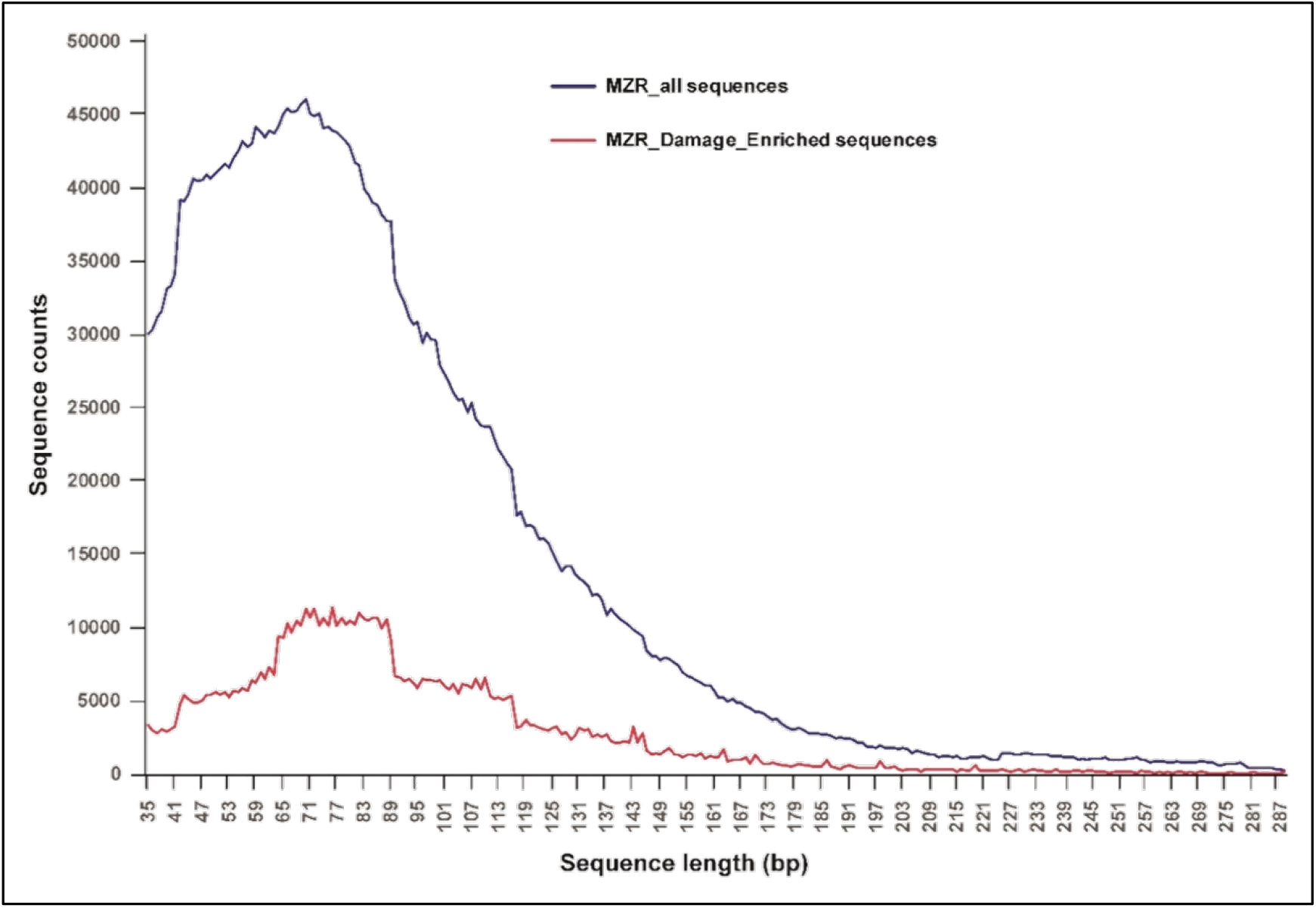
The sequence length distribution in MZR, including all sequences and the damage-enriched sequences, respectively. The average length of these two sequence sets are very similar (93bp vs. 92bp), supporting the authenticity of the MZR sequence data.

**Figure S4.**
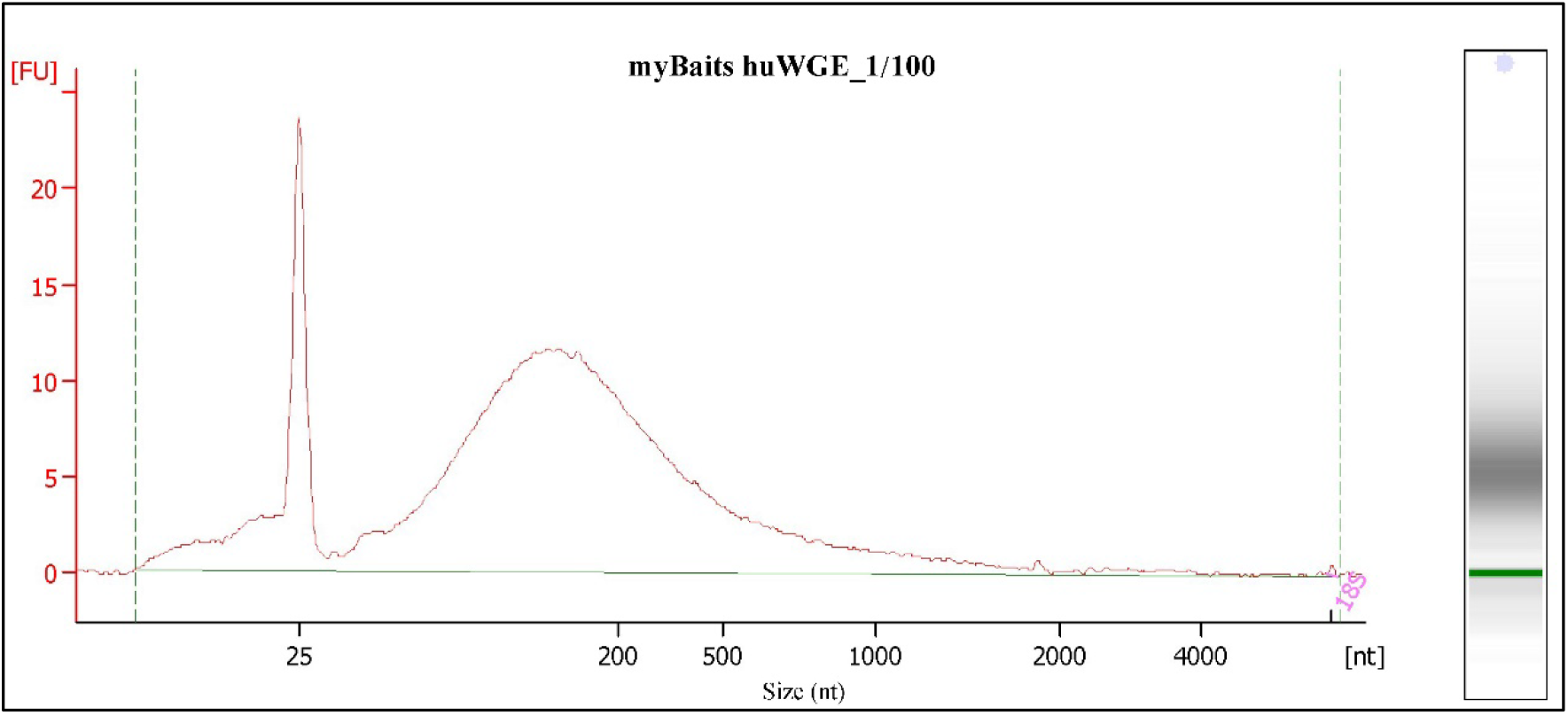
The bait length distribution for the MYbaits kit of MYcroarray, detected using the High Sensitivity DNA Kit of Agilent Bioanalyzer 2100. Before detection, we diluted the bait by 1/100.

**Figure S5.**
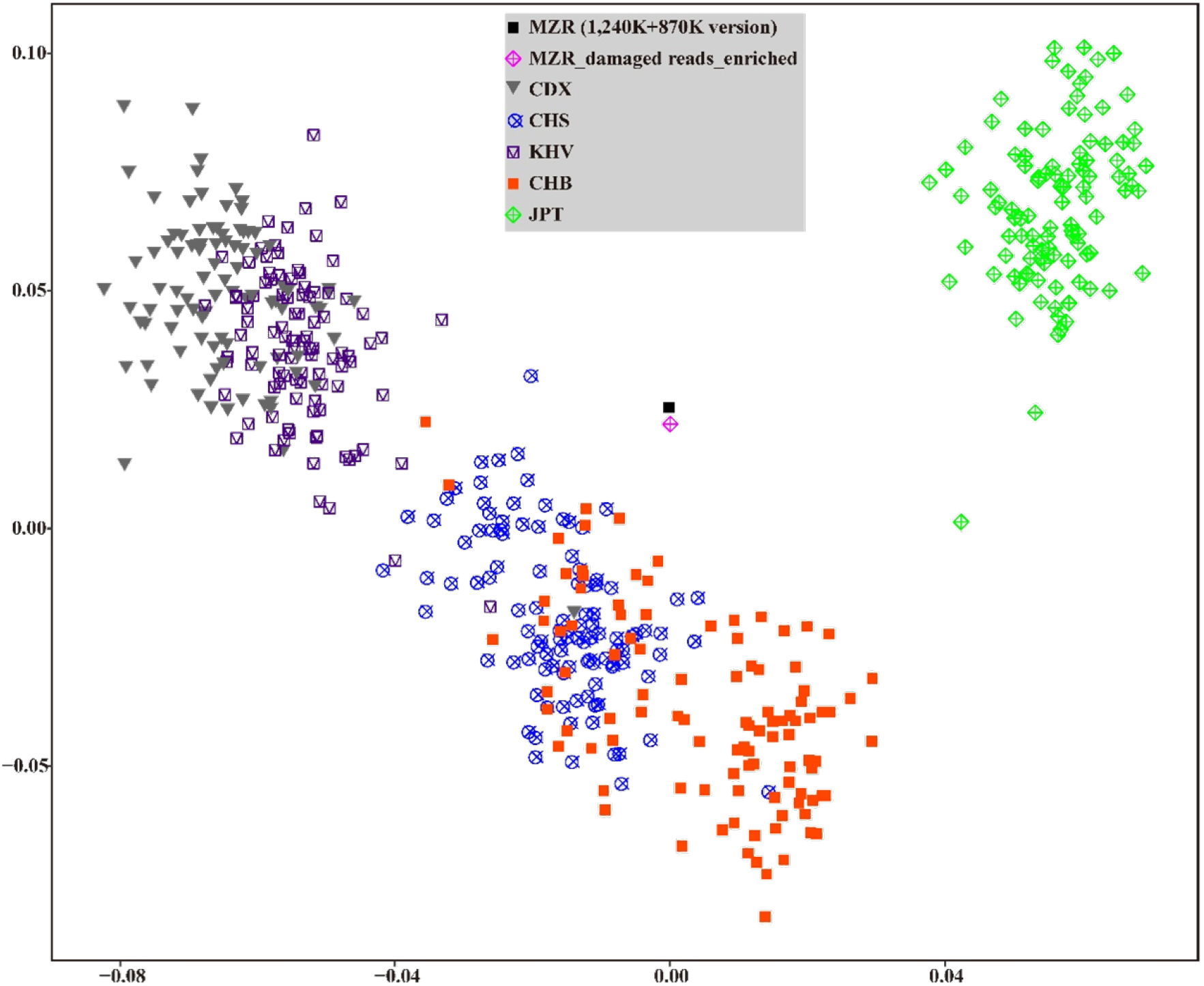
The PCA plot showing the relationship among MZR (using the original sequences and the damage enriched sequences) and present-day East Asians from the 1,000 Genomes Project^1^. We projected the MZR data onto the first and the second PC spaces established using the modern samples. We used the 1,240K+870K SNPs set for MZR data.

## Supplementary Methods

### Supplemental Analysis 1

We collected the East Asian mtDNA whole genome sequences from the literature ^2-65^, totally, we compiled 27,850 and 21,927 sequences of East Asians and Han Chinese. Using the *Haplogrep2* program (https://haplogrep.i-med.ac.at/), we obtained the mutation list for all the samples (Table S1), and summarized the frequency distribution of the 3,058 mismatch sites that Tabin *et al* observed (Figure 1A).

Furthermore, in order to strictly examine the MZR mtDNA mutation context and lineage definition, from the above damage enriched file, plus the all potential PCR duplicates removed file, we retrieved their mtDNA sequences and conducted a manual check using Integrative Genomics Viewer (IGV)^66^, and the mutation consistency contexts were summarized (Table S2), which are consistent with the result in our original paper.

### Supplemental Analysis 2

For the modern individuals, we selected East Asians from 1,000 Genome Project^1^, to retrieve the VCFs, and then we converted them into plink version files and merged with MZR data. For MZR data, by dividing them into two sets: the MZR-50% shortest sequences (MZR-short) and the MZR-50% longest sequences (MZR-long), using the *pileupCaller* program in *sequenceTools* (https://anaconda.org/bioconda/sequenceTools), we called the “pseudohaploid” genotypes, and generated 27,596 SNPs and 34,855 SNPs, respectively. For the 2.2 million SNPs (1,240K+870K) set, using the same method, we called the “pseudohaploid” genotype for MZR, MZR-short and MZR-long has 45,170 and 56,999 SNPs, respectively. We merged the above East Asian individuals from 1,000 Genome Project, the intersecting SNP number between the 1,240K+870K with 1,000 Genome dataset is 1,858,182. we performed PCA (using the --lsqproject parameter) to detect the difference between the two MZR datasets (Figure 1D).

In addition, with the sequence ends untrimmed mapped bam version, using *PMDtools*^67^, we enriched the damaged reads for MZR, and called 2.2 million SNPs set with the above method. It generated 12,177 SNPs when merging with the East Asians from 1,000 Genome Project, as well as the MZR all SNPs set, we performed PCA (using the --lsqproject parameter) to detect the difference between the two MZR datasets (Figure S5). We also plotted the sequence length distribution comparison between the all sequence set and damage enriched sequence set for MZR (Figure S3).

